# Fusogen-induced recovery of spinal cord function and morphology after complete transection

**DOI:** 10.1101/2025.09.13.674972

**Authors:** Michael Lebenstein-Gumovski, Tanzila Rasueva, Dmitriy Kovalev, Sergio Canavero, Alexander Zharchenko, Pavel Petrov, Andrey Zhirov, Andrey Grin’

## Abstract

**Background:** Spinal cord injury is a critical issue in neurosurgery, lacking established clinical methods for functional restoration. This study reports the effects of a fusogen sealant, composed of polyethylene glycol and chitosan, in an experimental model of complete spinal cord transection in pigs.

**Objective:** To evaluate the functional and morphological recovery of the spinal cord in an animal model of complete transection following treatment with a polyethylene glycol-chitosan conjugate.

**Materials and methods:** *Hungarian Mangalica* pigs (*m* = 20.0 *±* 2.0 kg, *N* = 5) underwent complete transection of the thoracic spinal cord, followed by an extended laminectomy and transpedicular fixation. In the experimental group (*N* = 3), a synthesized gel based on a polyethylene glycol-chitosan conjugate was applied to the spinal gap; the other group (*N* = 2) served as a control. The postoperative period lasted 60 days and included multi-component rehabilitation. Clinical-functional status was assessed using established neurological scales. *In vivo* retrograde tracing of the spinal cord was performed using hydroxystilbamidine (FluoroGold). Following the experiment, immunofluorescent histology was conducted using primary antibodies to neurofilament (NF-200), a fluorochrome-conjugated secondary antibody, and the nuclear dye 4’,6-diamidino-2-phenylindole (DAPI). The resulting morphology was examined via fluorescence and light microscopy.

**Results:** Control animals maintained lower paraplegia, anesthesia, and pelvic dysfunction throughout the experiment. In contrast, the experimental group showed positive changes, including the return of sensation from day two. By the end of the study, all animals in this group could assume an upright posture and ambulate on all limbs. These outcomes were statistically significant. Microscopy revealed axons traversing the injury site in the experimental group, whereas control samples showed degenerative post-traumatic changes.

**Conclusions:** This study demonstrates that a fusogen sealant based on a polyethylene glycol-chitosan conjugate promotes significant morphofunctional recovery after complete spinal cord transection, supporting its therapeutic potential.

## Introduction

Spinal cord injury with persistent neurological deficits is a disabling condition. No effective treatment currently exists to restore sensory and motor functions in affected patients. Approximately 27 million people worldwide live with a disability resulting from spinal cord injury [1]. Annually, the number of new cases reaches 8,000 in Russia, between 10,000 and 20,000 in the United States, up to 60,000 in China [2, 3]. These injuries are more common among the working-age population. In addition to drastically reducing a patient’s quality of life, spinal cord injuries impose a significant economic burden, making the search for an effective treatment method imperative.

Until recently, scientific efforts to address spinal cord injuries focused primarily on neural tissue regeneration. However, axonal regeneration in the spinal cord is hindered by a cascade of pathophysiological processes that lead to the formation of a complex pathomorphological structure, or glial scar, consisting of glial cells, fibroblasts, and connective tissue [4]. It is well established that the distal segment of transected axon undergoes Wallerian degeneration. Recent studies indicate that in mammals, the balance at the injury site shifts towards degeneration rather than regeneration [5].

Efforts to modulate these pathological processes have been largely unsuccessful for over a century. In recent decades, attention has turned to the mechanism of spontaneous axonal fusion observed in invertebrates, which allows for the highly specific restoration of injured axons without requiring regeneration. For instance, axon segments in nematodes can fuse, achieving *restitutio ad integrum* [6]. This spontaneous axonal repair mechanism is absent in mammals. Investigations into these processes led to the discovery of a new class of substances — fusogen sealants, such as polyethylene glycol (PEG) [7–10].

Our team synthesized a polyethylene glycol-chitosan conjugate (PEG-chitosan, Neuro-PEG) using an original methodology that proved effective in experiments involving rats and rabbits [11–14]. To improve clinical applicability, we developed a spinal cord restoration technique that combines a fusogen therapy protocol with controlled dorsal spinal traction, spinal cord segment resection, and rigid fixation [15]. Preliminary results demonstrated a positive effect of Neuro-PEG in pigs, using *in vivo* tracing with 1,1’-dioctadecyl-3,3,3’,3’-tetramethylindocarbocyanine (DiI) [16]. This article presents the results from an experiment on five *Hungarian Mangalica* pigs, that employed our method to assess its clinico-morphological effects via retrograde tracing.

Therefore, the objective of this study was to investigate the functional and morphological changes in the spinal cord after injury in an experimental model using a therapeutic approach that combines a Neuro-PEG, fusogen therapy, and rigid spinal fixation.

## Materials and methods

The study protocol was approved by the Local Ethics Committee of the Stavropol State Medical University (No. 97, April 15, 2021). This trial was carried out in accordance with the ethical standards of the European Convention for the Protection of Vertebrate Animals used for Experimental and other Scientific Purposes (European Treaty Series, No. 123, March 18, 1986) [17].

The experimental gel for intraoperative application was based on photo-crosslinked chitosan and covalently bonded PEG-chitosan, synthesized by our previously described method [12]. We used PEG (600 kDa; Merck, Germany) and chitosan (15 kDa; Merck, Germany) as starting materials. The PEG-chitosan conjugate was separated by centrifugation, washed twice with phosphate-buffered saline (PBS) and Type I water, then dissolved in PBS to 20 mg/mL and homogenized via ultrasonication.

### Surgery

Our surgical model involved female *Hungarian Mangalica* pigs (*m* = 20.0 *±* 2.0 kg, *N* = 5: 3 (experimental group), 2 (control group)). Three-month-old females were chosen because they allow easier urethral catheterization and monitoring of pelvic functions. Animals were anesthetized intramuscularly with a mixture of Zoletil (tiletamine/zolazepam; 3.0-7.0 mg/kg; Virbac, France) and Xylazine (2 mg/kg; Alfasan Int., Netherlands) and maintained on a propofol infusion (0.2 mg/kg/min; Propofol-Kabi, Fresenius Kabi, Germany) via a central venous catheter. After IV muscle relaxation with Rocuronium (1.2 mg/kg; B-Pharm, Russia) and catheterization, the animals were intubated and artificially ventilated with an air-oxygen mixture. A posterior approach was used to expose the laminae of the Th7–Th9 vertebrae, with dissection of the facet joints. An extended laminectomy with facetectomy of these segments created relative thoracic spinal instability. Transpedicular polyaxial screws (LLC Osteomed, Russia) were inserted at three levels (Fig 1a), and slightly curved longitudinal rods were positioned. Screw heads were approximated, dorsal traction was applied, and the rods were secured with washers (Fig 1b). Thereafter, a Smith-Petersen (chevron type) osteotomy and a pedicle subtraction osteotomy were performed to correct the fixed sagittal imbalance (Figs 1.1-1.6). For neuroprotection via local hypothermia, an ice slurry of frozen isotonic NaCl solution (*t* = 0 *±* 0.5 ℃) was placed on the dural sac. After 1 minute, a 2 cm wide area of the dural sac at T8 was cleared of the slurry, and the dura mater was opened and secured with holding threads. A flexible spatula was slipped under the spinal cord to protect the underlying dura, and the dentate ligaments were sectioned bilaterally. The spinal cord was then fully transected with a №11 scalpel. The resulting gap was irrigated with 2 mL of the PEG-chitosan gel (Figs 1c-1d). Simultaneously, a 50-mL intravenous drip infusion of PEG solution was administered. In control animals (*N* = 2), neither the PEG-chitosan gel nor the PEG infusion was applied. The cord segments were placed in the dural sac to achieve tight apposition. The dura mater was closed in a watertight manner and covered with adipose tissue (Fig 1e). Finally, the surgical wound was closed in layers over a vacuum aspiration drainage system.

**Fig 1.**
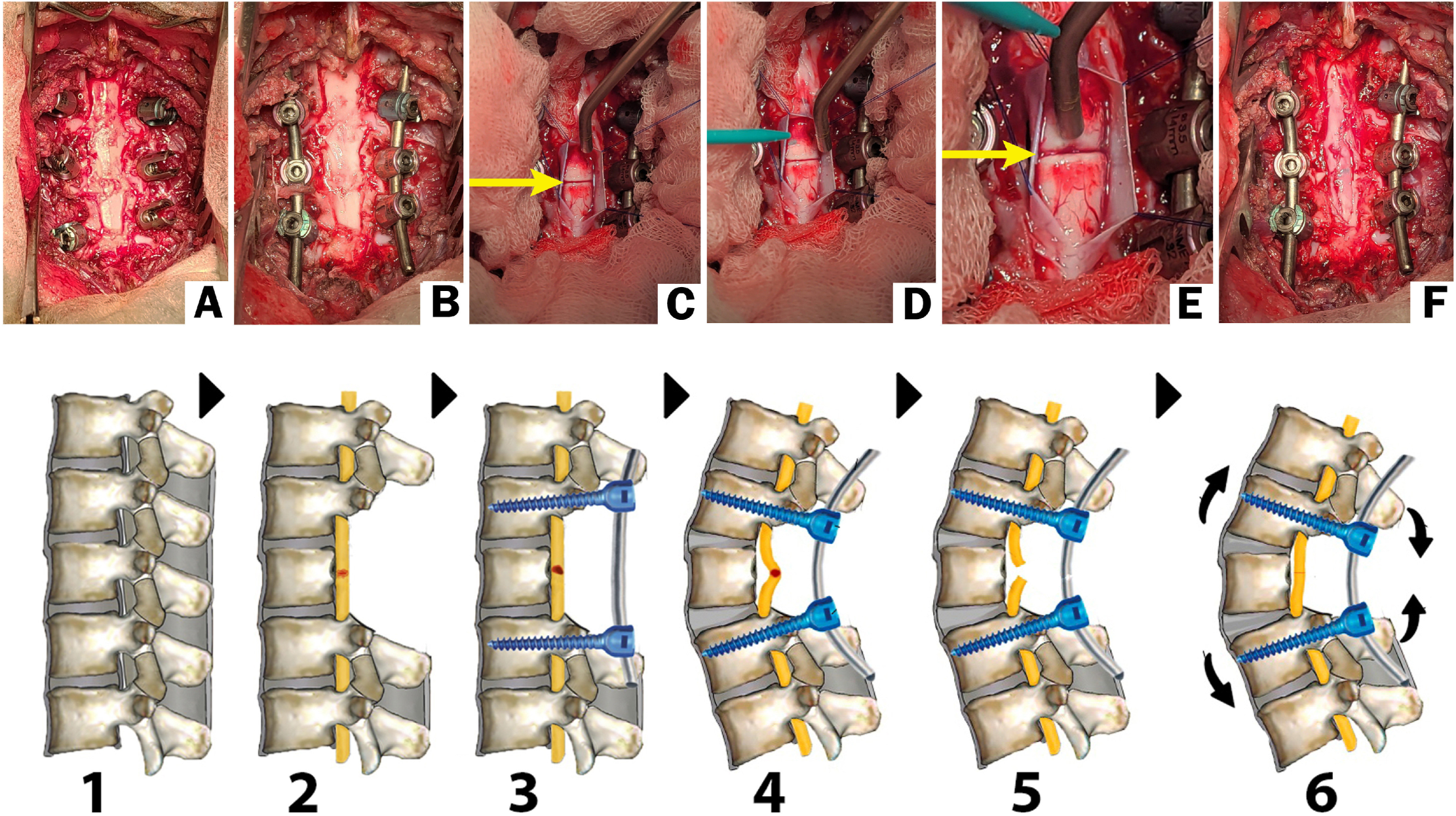
Stages of the surgical procedure. The top panel shows intraoperative photographs, and the bottom panel provides a corresponding schematic. Photographs of surgical stages: (A) Access to the spinal cord following laminectomy and installation of transpedicular screws; (B) Installation of longitudinal beams and application of traction to the screws; (C) Dissection of the dura mater, which is secured with holding threads before spinal cord transection; (D) Introduction of the PEG-chitosan gel into the transection gap. A close-up view of the transection site (yellow arrow) shows no bleeding after gel application; (E) The dura mater is sutured closed over the injury site; (F) Final appearance of the stabilized spinal fixation construct. For all photographs, the cranial end is at the top. Schematic of surgery: (1) Gaining surgical access; (2) Performing the laminectomy; (3) Installing transpedicular screws; (4) Applying traction to screw heads along the curved rod; (5) Transecting the spinal cord; (6) Performing the final fixation of screws.

### Postoperative care

The animals were extubated, transferred to spontaneous ventilation, and placed in recovery pens. In accordance with the European Convention for the Protection of Vertebrate Animals used for experiments or other scientific purposes [17], all animals received adequate analgesia (ketoprofen; 3 mg/kg; LLC Velpharm, Russia) and antibiotic therapy (intravenous cefoperazone sodium and sulbactam sodium; 25 mg/kg; Hebei Huachen Pharmaceutical, Co., Ltd., China). They were maintained on a controlled light-dark cycle with free access to water, regulated feeding, and environmental aconditioning at a maintained temperature. Daily massage of the hind limbs was performed to improve trophism and prevent pressure sores. Additionally, each limb underwent daily electromyostimulation (3–13 Hz frequency, 0.1–30 msec pulse duration) for 20 minutes twice a day. For 15 days postoperatively, animals received subcutaneous injections of neostigmine methylsulfate (18 µg/kg, twice daily) and intramuscular injections of trypsin (10 mg, once daily). For 7 days, animals in the experimental group received daily intravenous infusions of 50 mL of a 25% solution of PEG-600 (600 kDa; Merck, Germany) in 0.9% sodium chloride. During rehabilitation, animals were continuously encouraged to assume an upright position by being placed on all four limbs during feeding, with pelvic support. To monitor for central urinary dysfunction, the urethral catheter was removed every third day, and urine output was observed for 6 hours; if no voiding or signs of bladder filling were noted, catheterization was repeated. Defecation issues were managed with enemas.

### Neurological assessment

Neurological status was evaluated based on motor, sensory, and pelvic functions. Motor function was scored using the Individual Limb Motor Scale (ILMS) [18], the Pig Thoracic Spinal Cord Injury Behavior Scale (PTIBS) [19], and a modified Basso, Beattie, Bresnahan (BBB) motor activity scale [20], including sensory and pelvic function assessment [16]. For these evaluations, animals were released onto a 6-meter ramp with a soft surface, where their hind limb movements, gait, and any ataxia were recorded and scored. The return of nociception was tested using pinpricks. Ileus was evaluated via auscultation. The total follow-up period was 60 days.

### Statistical analysis

Statistical analysis of the data was performed using GraphPadPrism 10.5.0. A two-way analysis of variance (ANOVA) with a Tukey’s post hoc test was performed with significance set at P *<* 0.05. Pearson’s test was used for correlation analysis of the experimental group data between the behavioral scales.

### Histology

Fourteen days before the end of the experiment, one animal from the experimental group was anesthetized with Zoletil (see above) and given a series of microinjections of a FluoroGold solution (4%, 5 µL total, in 0.9% NaCl; Biotium, USA). The solution was injected into the spinal cord 5 cm below the injury site, using a Hamilton syringe over a 5-minute period via micro-access.

At the end of the 60-day follow-up period, the animals were euthanized with an overdose of Zoletil and Xylazine.

A 10-cm segment of the spinal cord containing the injury site was carefully isolated, removed, and immediately fixed for 24 hours in 4% paraformaldehyde in 0.1 M PBS at pH 7.4. For fluorescence microscopy specimens were cryoprotected in 20% sucrose at 4 ℃ for 48 hours and then sectioned into 30–40 µm thick longitudinal sections using a cryotome (Leica Microsystems CMS GmbH, Germany). For light microscopy, specimens were dehydrated through a graded series of alcohols (isopropanol) and embedded in paraffin using a Citadel 2000 carousel-type tissue processor (Thermo Fisher Scientific, USA). The resulting paraffin blocks were sectioned into 10-15 µm thick slices.

Sections were permeabilized by immersing slides in 0.3% Tween-20 (Abcam, UK) for 20 minutes. Non-specific binding was blocked by incubating slides in 1% fish gelatin (FGBA, Fish Gelatin Blocking Agent; Biotium, USA). Sections were then incubated for 1 hour at 25 ℃ with a mouse NF-200 monoclonal primary antibody (Abcam, UK) diluted in 1% FGBA. Following washes, slides were incubated in the dark for 1 hour with an Alexa Fluor 488-conjugated rat anti-mouse IgG2a monoclonal secondary antibody (Clone: SB84a; Abcam, UK). Cell nuclei were counterstained with a 300 nM solution of DAPI (Abcam, UK) in PBS. Finally, slides were air-dried and coverslipped with a mounting medium. The specificity of staining was assessed by excluding primary antibodies or incubating with a fish gelatine solution. The prepared paraffin sections were deparaffinized according to standard histological techniques. Subsequently, sections were stained with hematoxylin and eosin, with toluidine blue according to Nissl, and with Mallory’s polychrome stain to visualize connective tissue.

Fluorescence and light microscopy were performed using an AxioScope A1 microscope (Zeiss, Germany) and a BioTek Cytation-1 multimode cell imaging reader (Agilent Technologies BioTek, USA) with BioTek Gen5 software (Agilent Technologies BioTek, USA). Sections were examined and imaged, and the resulting photographs were processed using ImageJ software (NIH, USA).

## Results

### Neurological examination

Postoperatively, all animals initially exhibited lower paraplegia and anesthesia; these conditions persisted in the control group throughout the study. Pelvic functions were similarly impaired in all animals from both groups through postoperative day (POD) 2. A urethral catheter was maintained in all animals after surgery, with trial removals performed every 72 hours starting on day 3 for a 6-hour observation period. If urinary retention occurred, the catheter was reinserted. Constipation and urinary retention were pronounced in both groups during the POD 2, and fecal retention was managed with dietary adjustments, cleansing enemas, and abdominal massage.

In the experimental group, positive changes in motor activity and sensitivity were evident as early as POD 2. On that day, one animal demonstrated weak hind limb movements (a slight retraction when a limb was actively abducted by the experimenter), and all animals in the group responded to pinpricks at 3–4 of 6 tested sites on the hind limb skin. By POD 5, all animals in the experimental group had regained urinary control, allowing for catheter removal; in contrast, catheters in the control group were not removed until POD 10. On POD 7, one experimental animal made active attempts to stand using its hind limbs. Concurrently, the other animals exhibited sufficient hind limb strength to push off and resist the experimenter’s hand, though they did not yet attempt to stand. By POD 14, all animals treated with the PEG-chitosan conjugate displayed active hind limb movements and attempts to stand, albeit without weight-bearing support, along with a significant return of pain sensitivity. By POD 18, two experimental animals were capable of achieving a standing posture with support from their hind limbs.

By this stage, the animals were able to move independently using their hind limbs, though pronounced ataxia was evident. By week 4, their range of motion had increased considerably, and sensitivity scores were between 2 and 3, yet gait instability persisted. These parameters remained largely stable until the end of the study. On POD 60, all animals in the treatment group could move independently on all four limbs, demonstrated complete control over pelvic functions, and achieved a sensitivity score of 3.

This clinical progression is depicted in the photographs from the video storyboard of the gait analysis (Fig 2).

**Fig 2.**
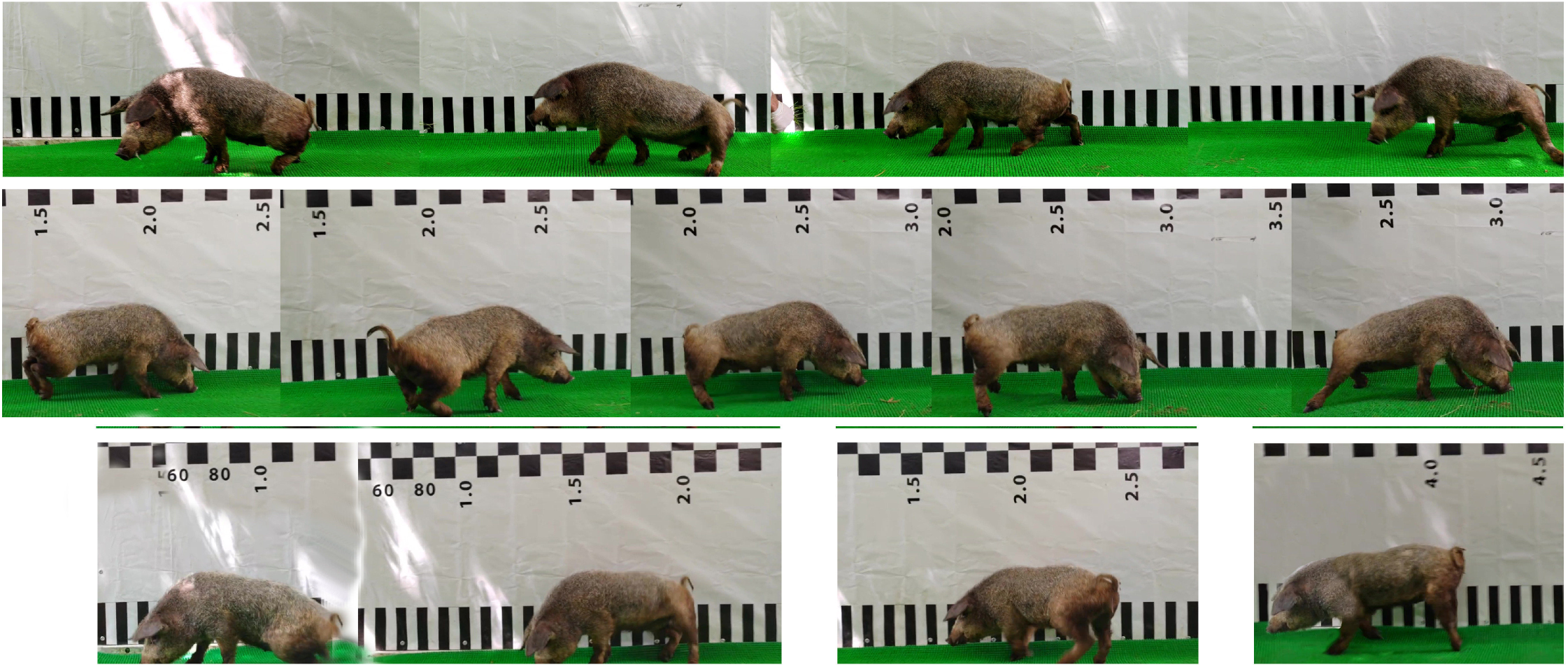
Progression of motor activity in a pig following complete spinal cord transection and application of the PEG-chitosan conjugate. The images are stills from video recordings at various POD. Top row (Day 10): The animal is noted to be making attempts to stand using its hind limbs. Middle row (Day 21): The animal can stand freely and attempts to walk. Bottom row (Day 40): The range of motion has increased significantly, though gait instability persists.

The line charts in Fig. 3 plot the average neurological scores for the experimental and control groups during the 60-day experiment. The trend line for the control group plateaued at a minimum level, indicating no functional recovery. In contrast, the trend line for the experimental group shows a steady improvement in scores, reflecting the recovery of sensory, motor, and pelvic functions.

**Fig 3.**
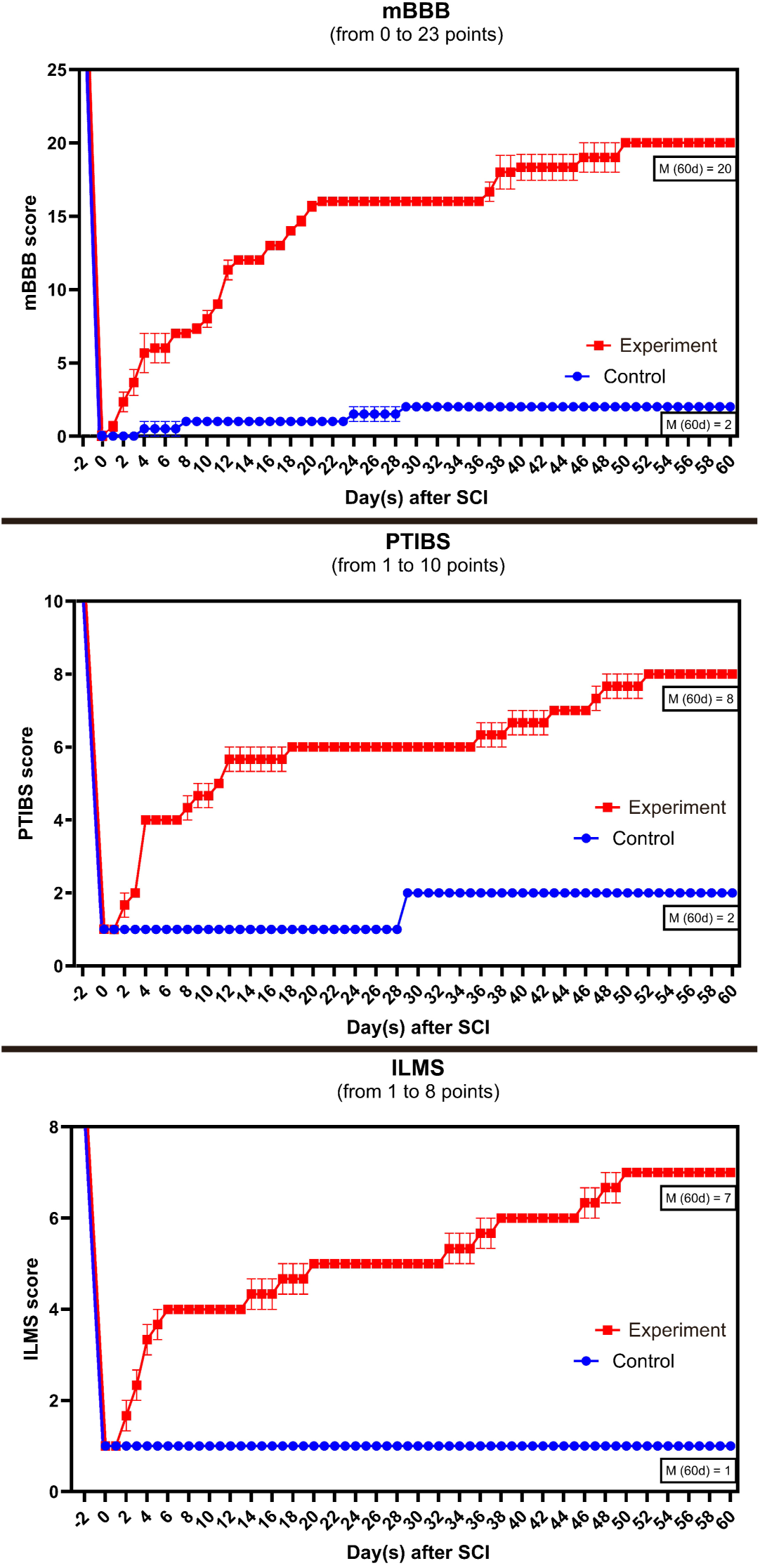
Neurological scores over a 60-day period for the experimental and control groups. The assessment was made using three scales: the modified Basso, Beattie, Bresnahan (mBBB) scale, which includes sensory and pelvic function criteria; the Pig Thoracic Spinal Cord Injury Behavior Scale (PTIBS); and the Individual Limb Motor Scale (ILMS). Line charts display the progression of individual animals throughout the experiment. Data are presented graphically using *M ±* SEM (standard error of the mean). The graphs show the mean (*M*) at 60 days after surgery.

Using the Tukey’s test, statistical analysis of data showed a significant difference (p *<* 0.001) on all scales between the control and experimental groups over a 60-day period. The bar charts (Fig 4), which display the mean neurological scores, confirmed these differences were highly significant. Correlation analysis of the data revealed a high level of correlation between the behavioral scales using the Pearson criterion (r from 0.95 to 1.0) for experimental group (Fig 4).

**Fig 4.**
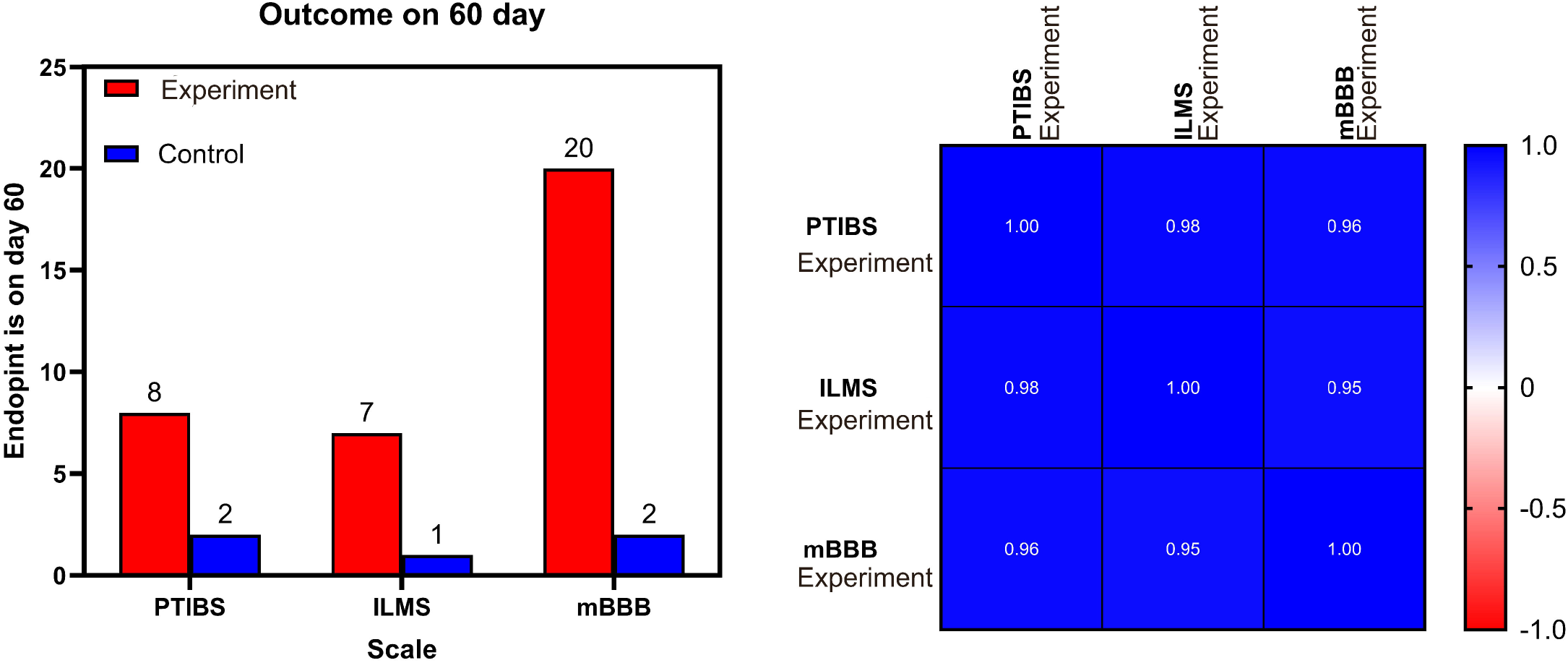
Results of statistical analysis of experimental data. Bar charts of mean scores for the three neurological scales used in the POD 60 (on the left). Using Pearson’s criterion for correlation analysis of experimental group data between behavioral scales (on the right).

### Microscopy

In the control group, histological analysis revealed the transection site. The sections showed an absence of axons in the immediate injury area and reduced axonal density beyond it, which is indicative of Wallerian degeneration. A corresponding decrease in neurofilament content was noted in the caudal segment of the spinal cord, further signifying Wallerian degeneration. In the injury zone, tissues stained intensely with DAPI but lacked neurofilaments, a finding that reflects glial proliferation and scarring (Fig 5). As DAPI is a nuclear stain and neuron-specific markers like NF-200 are absent in the gap (diastasis), this dense cellular staining is interpreted as glial cell proliferation filling the injury site.

**Fig 5.**
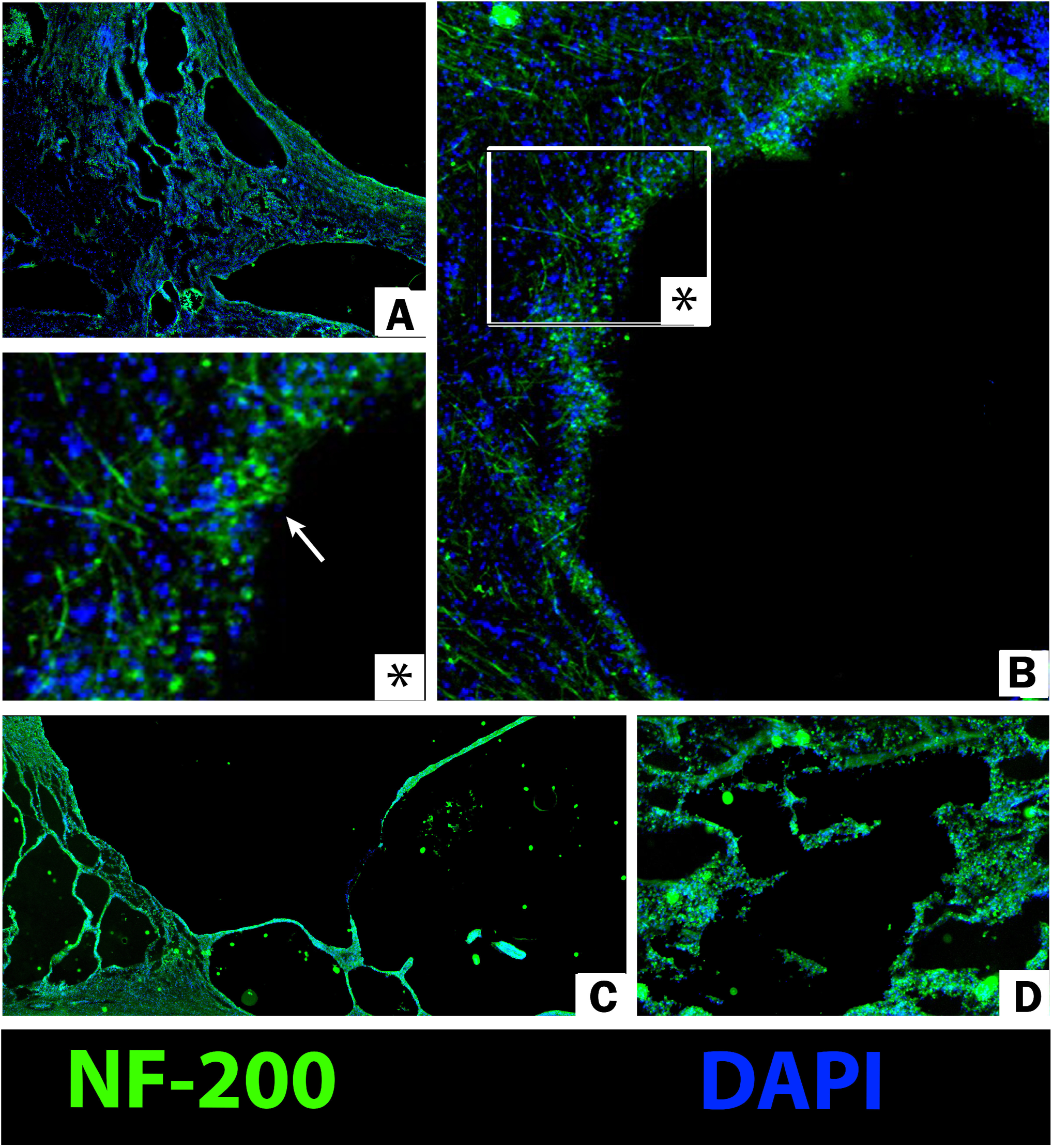
Immunofluorescence histology of the spinal cord from the control group. The images show cyst formation in the area of the spinal cord injury, a predominance of glial cells, and no growth of axonal structures through the cyst. (A) View of multiple cysts. (B) A cyst cavity with axonal retraction bulbs; the asterisk (*) indicates an enlarged area of the cyst wall. (C) An extensive cyst corresponding to a decreased NF-200 signal. (D) Disorganized, loose axonal structures with some background non-specific staining. Sections were stained with an antibody for NF-200 and the nuclear stain DAPI. Magnifications: A, D - ×40; C - ×25; B - ×100; * - ×400.

In samples from the experimental group, the injury site exhibited an increased density of neurofilaments. Although isolated cysts were noted in the gray matter, the white matter displayed twisted, thickened axons traversing the lesion area (Fig 6).

**Fig 6.**
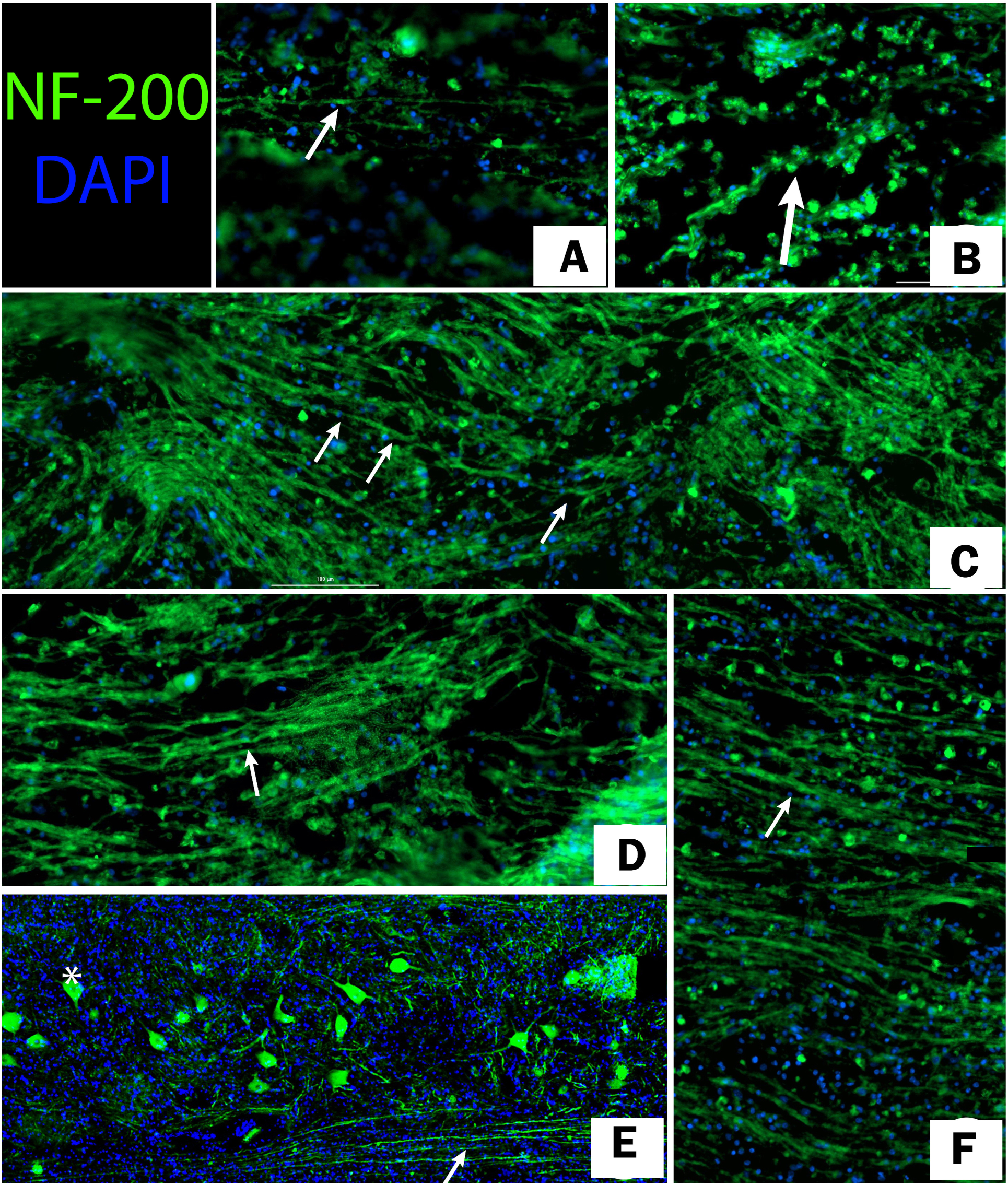
Immunofluorescence histology of the spinal cord from the experimental group. The horizontal section displays the proximal end of the spinal cord (left), the injury area (center), and the distal segment (right), with the cranial end to the left. (A-F) Arrows indicate axons crossing the injury site. (E) A gray matter area with neurons adjacent to the injury. The asterisk (*) indicates the body of a neuron in the gray matter. The image shows the merged channels for NF-200 and nuclear DAPI staining. A, B - ×400; C, E, F - ×100; D - ×200.

In the animal subjected to retrograde axonal tracing, FluoroGold dye was observed to have spread along axons that traversed the injury site (Fig 7 and S1 Fig). The density of these labeled axons appeared lower within the injury area compared to the distal spinal cord, a finding attributable to both the selective spread of the dye along only the traced axons and to partial axonal degeneration at the injury site itself.

**Fig 7.**
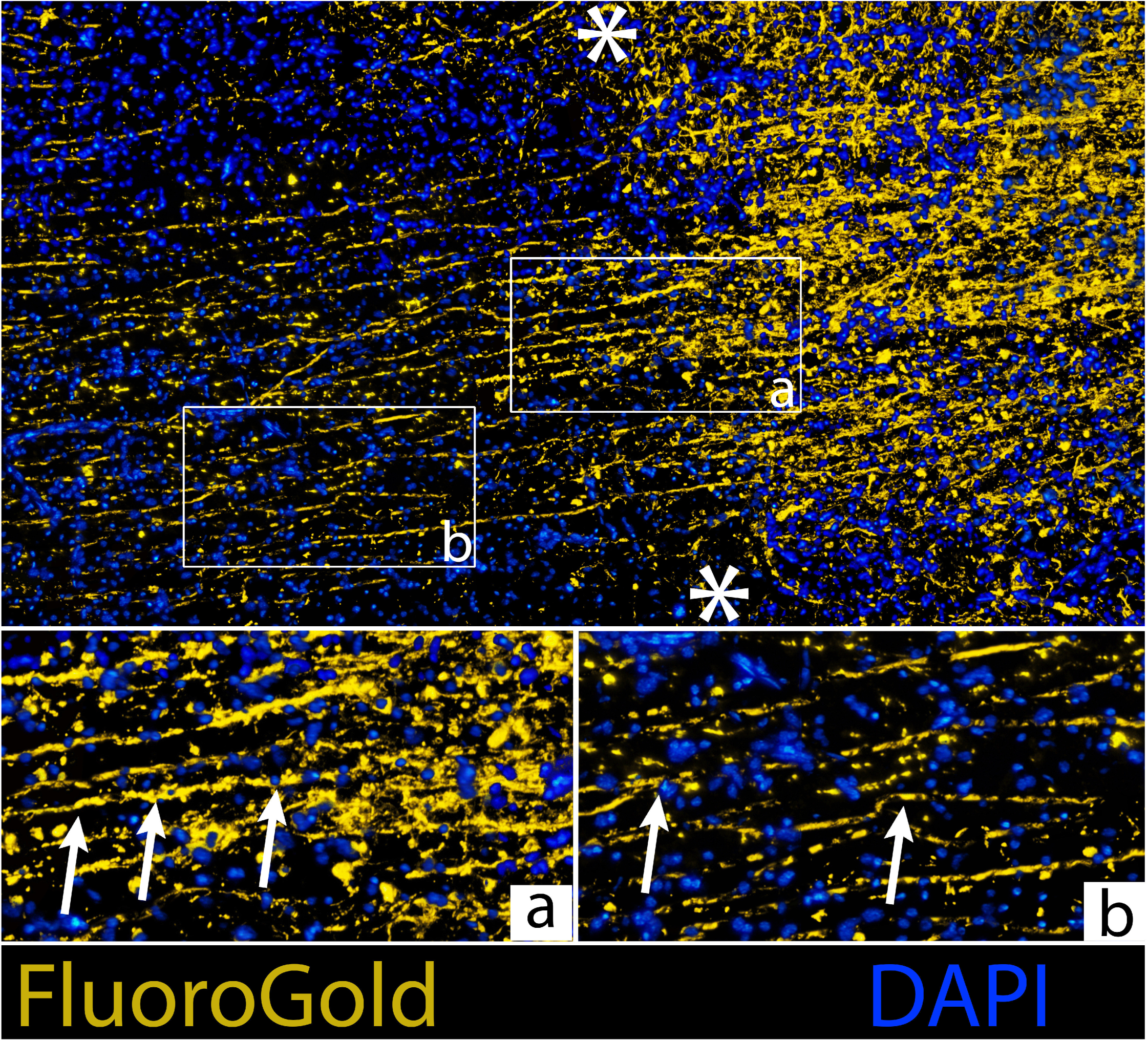
*In vivo* tracing of spinal cord axons in the experimental group. The images are horizontal sections with the cranial end on the left. The ends of the injury site are marked by asterisks. Nuclear staining with DAPI (blue), axons are traced with tracer FluoroGold (yellow). The superposition of channels is shown above (×100). Arrows indicate axons crossing the injury site at fields a and b (×400).

Light microscopy of samples from the control animals revealed gross degenerative processes affecting the nerve tissue (Fig 8). The analysis showed that the primary injury, a smooth transverse cut of the spinal cord, led to pathological changes that extended far beyond the immediate transection site.

**Fig 8.**
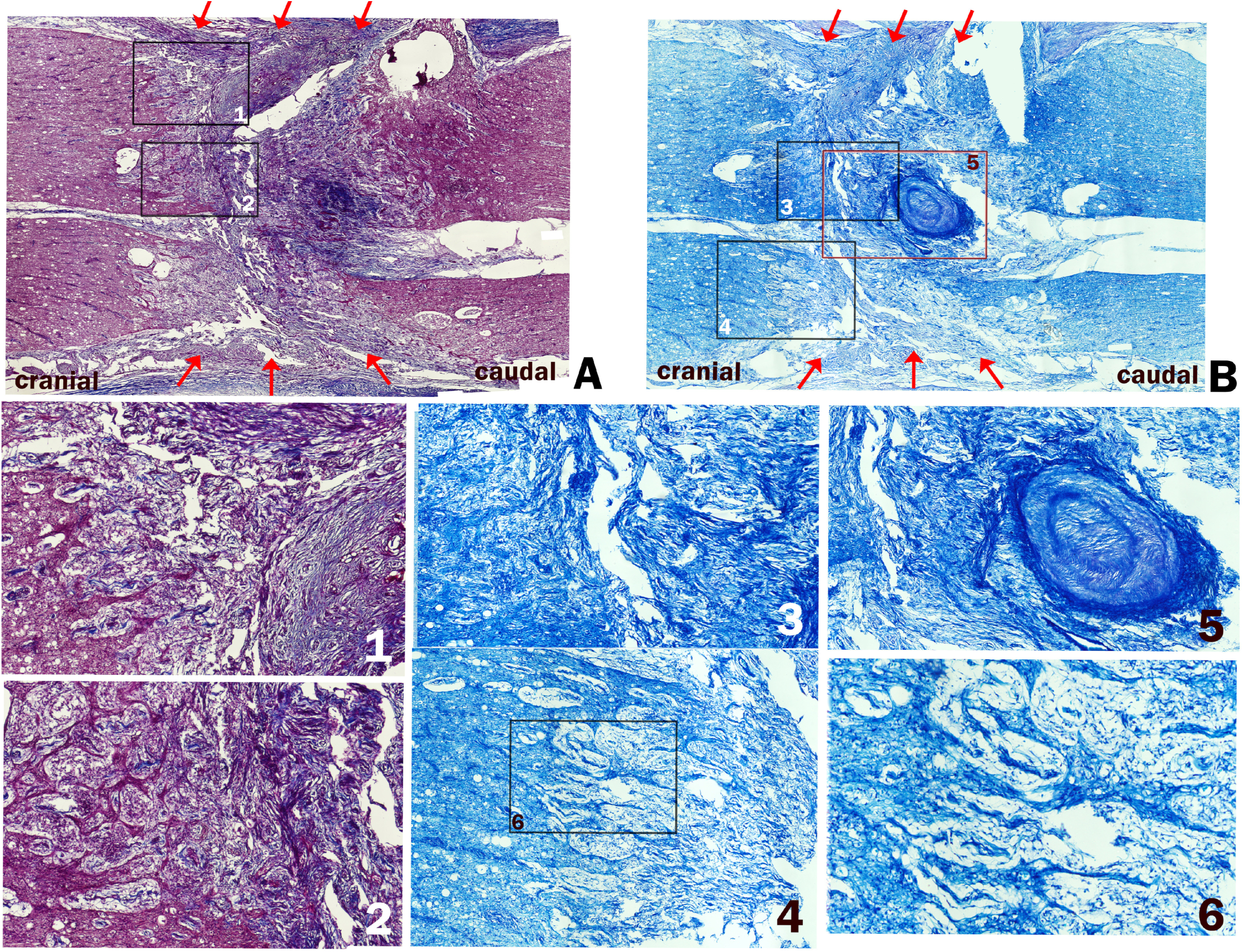
Pronounced degenerative changes at the spinal cord injury site in a control animal showed by light microscopy. The cranial end is to the left, and arrows indicate the transection site. (A) Section stained with Mallory’s trichrome (×200). (B) Section stained with toluidine blue according to Nissl (×200). Numbered labels indicate areas of: (1) nerve tissue degeneration and a fibro-glial scar in the proximal segment; (2) a fibro-glial scar with chaotically arranged axonal stumps and club-shaped thickenings; (3) fibrous changes within the transection gap (diastasis); (4) swelling and degeneration of proximal axon segments; (5) connective tissue organization of necrotic tissue; and (6) degeneration and swelling of axons.

Multiple hemorrhages, periaxonal edema, glial edema, glial proliferation, infiltration by erythrocytes and macrophages, decay of nervous tissue with the formation of necrotic detritus undergoing replacement by connective tissue were found in the distal and proximal sections of the spinal cord, extending 1-2 segments above and below the transection site (Figs 8 and 9).

**Fig 9.**
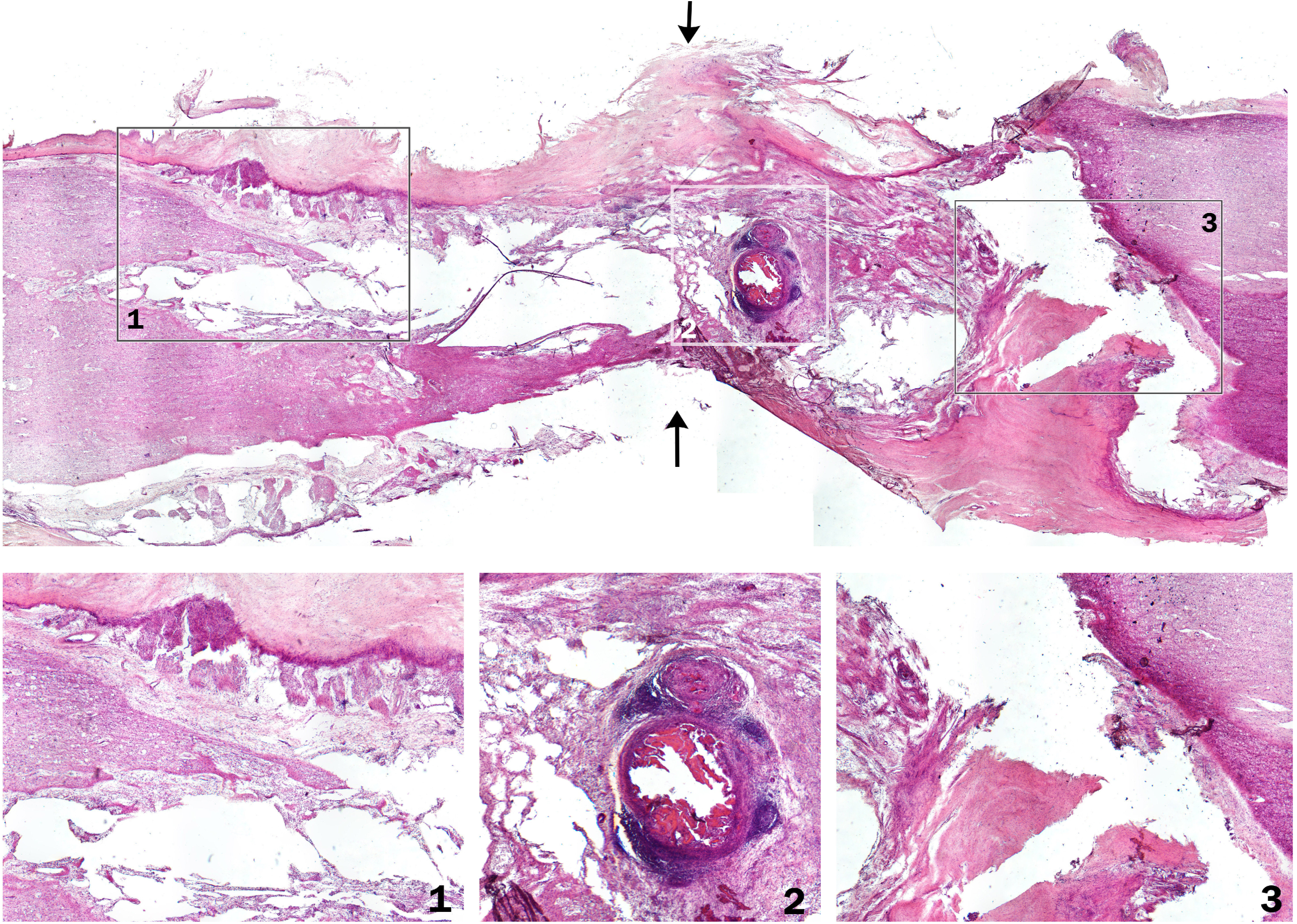
General view of the injured spinal cord from a control animal, stained with hematoxylin and eosin (×400). The image shows prominent degenerative changes, with cystic cavities affecting several spinal segments. The cranial end is to the left, and arrows indicate the transection site. Numbered labels indicate: (1) an area of nerve tissue degeneration and a fibro-glial scar in the proximal segment; (2) the central injury area with cysts, hemorrhages, and necrotic changes; and (3) degenerative changes in the distal segment.

In the experimental group, tissue samples displayed smaller areas of secondary damage (Fig 10). The extent and number of cysts, hemorrhages, and necrotic foci were reduced, and the area of altered tissue was mainly confined to the primary injury site. Notably, the injury site was crossed by a significant number of altered, twisted, and thickened axons, forming “axonal bridges” that connected the distal and proximal sections of the spinal cord.

**Fig 10.**
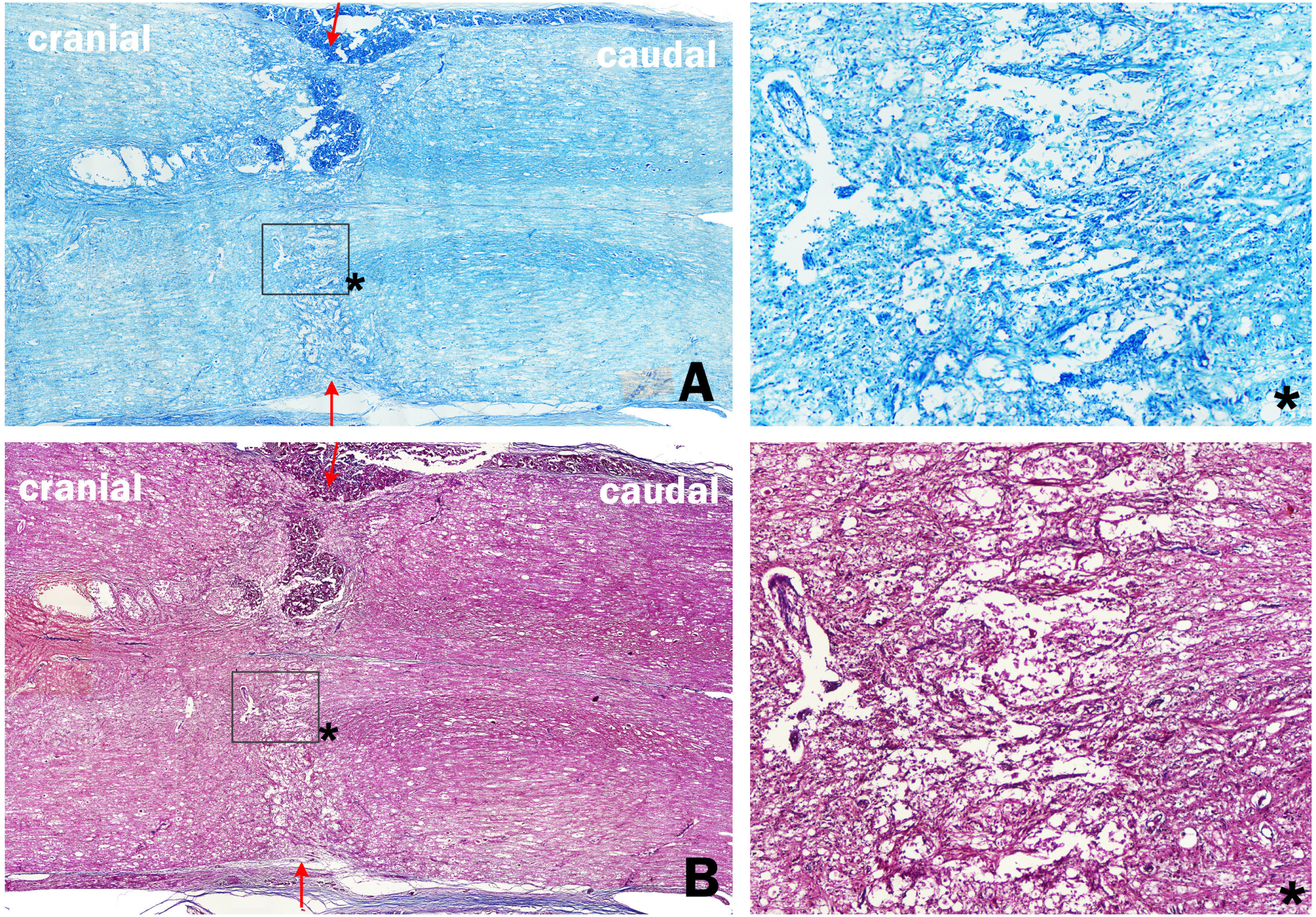
General view of the spinal cord from the experimental (PEG-chitosan) group. Less pronounced degenerative changes and the presence of “axonal bridges” in the injury area are shown. The cranial end is to the left, and arrows indicate the transection site. (A) Section with Nissl staining using toluidine blue (×200). (B) Section with Mallory’s stain (×200). Asterisks indicate regions that are shown at higher magnification in the corresponding insets on the right.

## Discussion

Until recently, experimental approaches to treating spinal cord injury focused on facilitating nervous tissue repair and creating conditions to support its regeneration [5, 21, 22]. *In vitro*, mammalian central nervous system axons demonstrate regenerative capabilities similar to those of peripheral nerve axons upon injury [21].

However, due to the unique cellular environment of the central nervous system, robust axonal regeneration in vertebrates is primarily observed in amphibians and fish, and to a lesser extent, in birds. Although mammals retain some regenerative potential, it is extremely limited. Following a spinal cord injury, a cascade of pathological reactions ensues, creating a vicious cycle that leads to the extensive death of surrounding healthy cells. This secondary injury cascade, which includes processes such as apoptosis, excitotoxicity, the release of prostaglandins, interleukins, and tumor necrosis factors, as well as the activity of fibroblasts and macrophages, overwhelms intrinsic regenerative mechanisms and renders axonal repair largely ineffective.

Alternatively, mechanisms more energetically favorable than regeneration exist for restoring spinal cord function [6, 23–26]. In some lower vertebrates, endogenous fusogen molecules are known to facilitate the fusion of damaged axon membranes. This fusion mechanism is highly specific to each axon and functional group, allowing the proximal and distal segments of a severed axon to “find” each other and reconnect across the gap [6]. Research indicates that it is possible to trigger a similar recovery mechanism in mammals by using exogenous fusogen sealants [7–10].

Given the rapid clinical improvement observed, the therapeutic effects of the PEG-chitosan conjugate synthesized by our team cannot be attributed solely to axonal regeneration, which is a much slower process. This points to immediate neurorepair mechanisms namely axonal fusion, being the primary driver of the initial recovery. We expect that, in the long term, neuroregenerative processes could also occur, further enhancing the clinical outcome.

However, the positive outcome is likely attributable not only to the experimental gel but also to the entire suite of measures applied in the study. For the effective use of PEG-chitosan, certain conditions must be met. Tight apposition of the spinal cord stumps requires smooth, even surfaces, which in a clinical scenario may entail resecting the trauma zone and performing a shortening spondylodesis or subtraction spondylotomy. Transpedicular fixation stabilizes the vertebral column and ensures the proper alignment of the spinal cord segments. Local cooling provides neuroprotection by slowing oxidative metabolism, preventing hypoxia and delaying the cytotoxic cascade. Since the intraoperative application of the fusogen is brief, with the initial repair process lasting only seconds to hours, postoperative intravenous infusions are important for maintaining a therapeutic concentration of the fusogen at the injury site. Finally, rehabilitation measures including massage, motivation, and electrical stimulation further enhance the treatment’s effectiveness. These results confirm the reproducibility of our method, consistent with our previous study in pigs that yielded similar outcomes [16]. Conducting this experiment in pigs, a highly relevant translational model from both physiological and anatomical perspectives, marks a significant step forward in the preclinical development of this method.

## Conclusion

This study demonstrates that fusogen-induced axonal repair is achievable after an experimental spinal cord injury. The use of a PEG-chitosan fusogen sealant, combined with subsequent postoperative intravenous PEG infusions, resulted in functional recovery in the treated animals, with motor and sensory outcomes directly correlating with the application of the therapy. Immunofluorescence with DAPI and NF-200 antibodies, staining and FluoroGold tracing revealed numerous axons crossing the injury site in the treated pigs. In contrast, light microscopy of control animals showed classic degenerative processes, including periaxonal and pericellular edema, as well as extensive foci of necrosis and fibrosis that spread far beyond the transection site. The histological results from the experimental group, which showed fewer degenerative changes and the presence of “axonal bridges” crossing the injury site, indicate that the methods used confer not only direct axonal repair but also a neuroprotective effect.

Thus, this study indicates that fusogen-induced spinal cord repair is possible and that fusogen therapy may become an effective treatment method for spinal cord injury in the near future. It is worth noting, however, that the fusogen alone is likely insufficient to achieve this effect. A combination of several factors, such as the precision of the surgical cut, neuroprotective strategies like local hypothermia, and a supportive postoperative environment, is probably required to create the optimal conditions for fusogenic action and successful axonal fusion.

However, for the clinical translation of these findings, it is critical to first understand the fundamental mechanisms of fusogenic nervous tissue repair. Elucidating these mechanisms should be a primary focus of future research.

## Supporting information

**S1 Fig. *In vivo* tracing of spinal cord axons at the site of transection in the experimental group.** The images are horizontal sections with the cranial end on the left. Nuclear staining with DAPI (blue), axons are traced with tracer FluoroGold (yellow) (×100).

## Acknowledgments

We thank Andrey Panferov, Alina Nurislamova for management and language support of this article.

## Author Contributions

**Conceptualization:** M.V. Lebenstein-Gumovski.

**Data curation:** M.V. Lebenstein-Gumovski.

**Formal analysis:** T.S.-M. Rasueva, A.V. Zharchenko.

**Investigation:** M.V. Lebenstein-Gumovski, T.S.-M. Rasueva, D.A. Kovalev, A.V. Zharchenko, P.O. Petrov, A.M. Zhirov.

**Methodology:** M.V. Lebenstein-Gumovski, A.V. Zharchenko.

**Project administration:** M.V. Lebenstein-Gumovski.

**Supervision:** M.V. Lebenstein-Gumovski.

**Writing – original draft:** M.V. Lebenstein-Gumovski, T.S.-M. Rasueva, D.A. Kovalev.

**Writing – review & editing:** M.V. Lebenstein-Gumovski, S. Canavero, A.A. Grin’.

## Ethical Approval

The study protocol was approved by the Local Ethics Committee of Stavropol State Medical University on April 15, 2021. The study was conducted in accordance with the ethical standards for the treatment of animals established by the European Convention for the Protection of Vertebrate Animals used for Experimental and Other Scientific Purposes.

## Use of artificial intelligence (AI)

The authors confirm that there was no use of artificial intelligence (AI)-assisted technology for assisting in the writing or editing of the manuscript and no images were manipulated using AI.

