## Supplementary figures and images for "Fusogen-induced recovery of spinal cord function and morphology after complete transection"

### S1 Fig

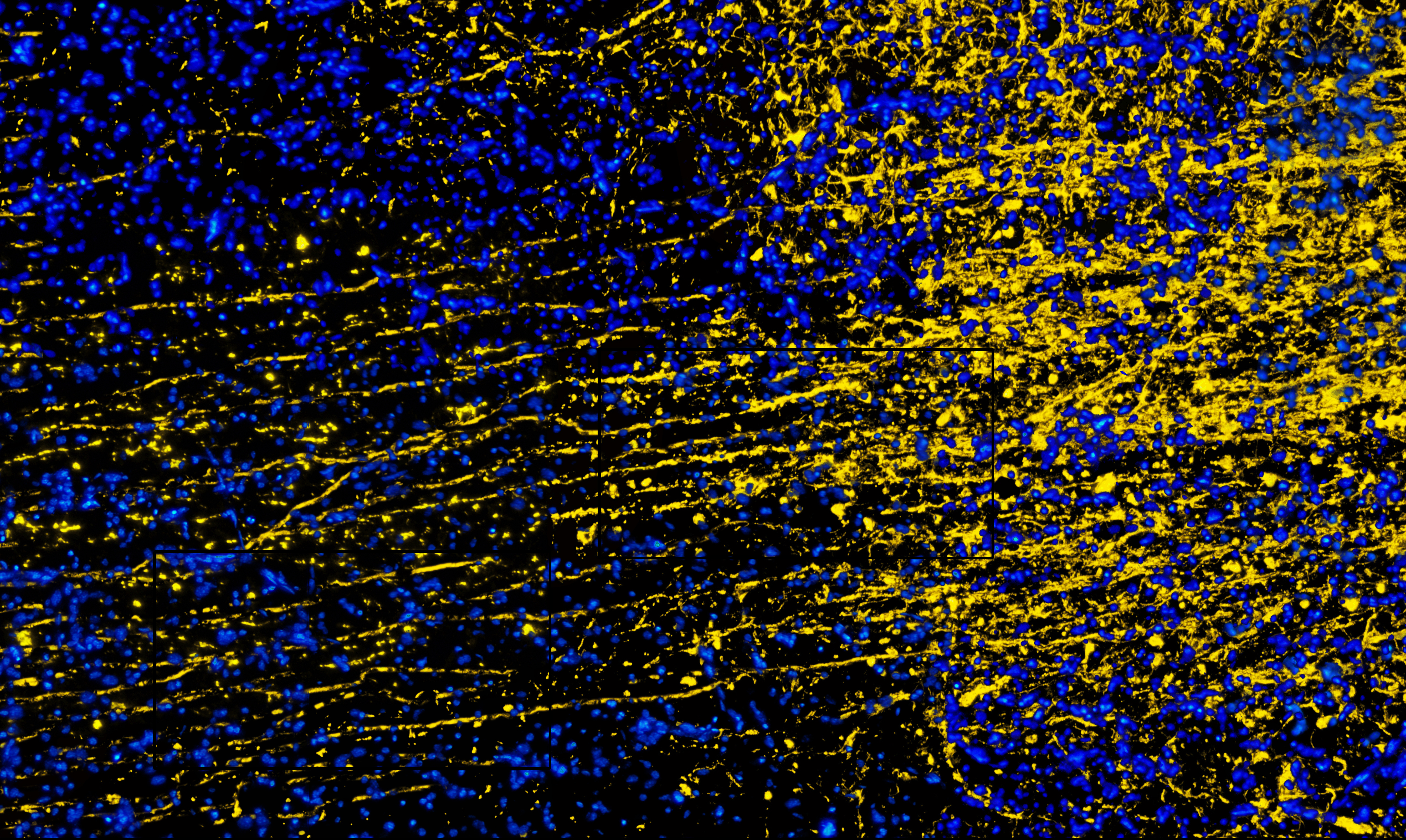
